# MkAtt-SDN2GO: Multi-kernel Attentive-SDN2GO Network for Protein Function Prediction in Humans

**DOI:** 10.1101/2025.11.02.686093

**Authors:** Kartik Jhawar, Tapasvi Bhatt, Rohan Sunil, Wang Lipo

## Abstract

Accurately annotating the functions of uncharacterised human proteins remains a major bottleneck in biology. We present **MkAtt–SDN2GO**, a neural architecture that extends SDN2GO by integrating adaptive multi-kernel convolution and attention mechanisms to predict Gene Ontology terms from protein sequences, domains, and protein–protein interaction (PPI) context. The sequence stream employs a learnable multi-kernel convolution layer that combines features from multiple kernel sizes through attention-based gating, enabling adaptive motif detection without relying on a fixed receptive field. A self-attention layer models long-range dependencies, while cross-attention integrates sequence, domain, and PPI representations into a unified prediction space. On a CAFA-style benchmark, MkAtt-SDN2GO improves Molecular Function (MF) F_max_ by 14.8% (0.657 vs 0.572) and Recall_max_ by 18.8% over SDN2GO. Across *Homo sapiens*, the fused model achieves top F_max_ scores in Biological Process (BP) (0.441), MF (0.657), and Cellular Component (CC) (0.522) compared with other methods. Although the domain-only stream performs strongly, the cross-attention fusion enhances robustness and interpretability when individual modalities are weak or missing. Overall, adaptive multi-scale convolution combined with attention thus advances large-scale protein annotation and offers a scalable and potential tool for functional genomics and disease research.

## Introduction

The sequencing of the human genome marked a monumental achievement in biology, yet a significant challenge remains: a substantial portion of human proteins lack functional characterization. These “uncharacterized” proteins represent critical gaps in our understanding of fundamental cellular processes, disease mechanisms, and potential therapeutic targets. While traditional experimental methods for determining protein function are invaluable, they are often laborious and resource-intensive. This necessitates the development and application of robust computational approaches capable of predicting protein functions at scale, thereby generating testable hypotheses and accelerating biological discovery.

Protein–protein interaction (PPI) networks offer a powerful framework for computational function prediction. These networks map the complex web of physical associations between proteins within a cell, providing insights into cellular organization and function. The principle of “guilt by association” underlies many network-based approaches: proteins that interact or reside within the same network module often share related biological roles. Early efforts, such as the work by Goehler et al. [2004] on Huntington’s disease, demonstrated the potential of PPI networks derived from yeast two-hybrid screens to identify novel interactions and assign functions to previously uncharacterized proteins. Subsequent large-scale studies, like that of Stelzl et al. [2005], expanded these efforts, generating vast human PPI networks and developing scoring systems to enhance prediction confidence, although highlighting the persistent challenge of false positives inherent in high-throughput experimental data.

Analyzing the topology of PPI networks, particularly through clustering algorithms, has proven effective in identifying functional modules – groups of proteins collaborating in specific biological processes. Silva et al. [2017], for instance, constructed a testis/sperm-specific network and used clustering to implicate uncharacterized proteins in spermatogenesis. Similarly, Nuncia-Cantarero et al. [2018] combined PPI network analysis with transcriptomics to identify functionally relevant gene clusters in breast cancer. These studies underscore the utility of network structure in revealing biological organization, though interpreting the functional significance of identified clusters requires careful integration with other data types. Multi-scale clustering approaches, employing algorithms like Markov Clustering (MCL) Van Dongen [2000] for fine-grained complexes and Louvain Blondel et al. [2008] for broader modules, offer a more comprehensive view by capturing functional organization at different resolutions.

The increasing availability of sequence and interaction data has spurred the development of sophisticated computational and machine learning methods for function prediction. Tools like PANNZER Koskinen et al. [2015] employed statistical methods like weighted k-nearest neighbors to predict GO terms and free-text descriptions with high throughput. More recently, deep learning has emerged as a particularly promising direction. One notable framework is SDN2GO Cai et al. [2020], which proposed an integrated deep-learning classification model to predict GO terms. The SDN2GO model specifically applied convolutional neural networks (CNNs) to learn and extract features independently from protein sequences, protein domains, and known PPI networks. A key aspect of their approach involved processing domain information using techniques inspired by Natural Language Processing (NLP) and utilizing a pre-trained deep learning sub-model to capture comprehensive domain features. These diverse features extracted by the CNNs were then integrated using a specialized weight classifier to achieve accurate GO term predictions. SDN2GO was rigorously evaluated using CAFA-style time-delayed datasets and demonstrated superior performance compared to baseline methods like BLAST and other contemporary approaches at the time.

Building upon these foundations, recent advancements have focused on incorporating attention mechanisms, inspired by their success in natural language processing Vaswani et al. [2017]. Attention allows models to dynamically weigh the importance of different parts of the input data when making predictions. For instance, PFresGO Pan et al. [2023] utilizes self-attention to model relationships between GO terms and cross-attention to align protein sequence embeddings with GO term embeddings. HNetGO Zhang et al. [2023] integrates heterogeneous network data using attention-based graph neural networks. These attention-based models, including others like Chen et al. [2022] and Wang et al. [2023], have demonstrated improved performance, particularly in capturing complex dependencies and long-range interactions within sequences and networks, often leading to better prediction accuracy, especially for less frequent GO terms Gligorijevic et al. [2021]. Hybrid approaches combining structure prediction, homology, and network inference Zhang et al. [2018], Duek and Lane [2019] further enrich the landscape of prediction strategies. In contrast to these prior works, none have explicitly combined an adaptively weighted multi-kernel convolutional layer with a self-attention mechanism for protein function prediction. Several prior methods incorporate structural or shape-related cues through auxiliary modules–for instance, by exploiting predicted three-dimensional structures or low-resolution shape proxies Zhang et al. [2018], Duek and Lane [2019], Gligorijevic et al. [2021], or by scanning for sequence motifs with fixed-size convolutional filters Kulmanov and Hoehndorf [2020]. Size-adaptive, learnable kernels have also been proposed to adjust the receptive field on the fly Tek et al. [2021], Li et al. [2019]. Although these strategies introduce valuable geometric priors, they are chiefly designed for semantic-segmentation accuracy or robustness enhancement and seldom provide particle-level interpretability; moreover, they rarely couple local features with explicit relational modeling. Attention-based approaches, on the other hand, emphasize feature re-weighting and context aggregation Pan et al. [2023], Zhang et al. [2023], Chen et al. [2022], Wang et al. [2023], but typically keep convolution and attention as separate stages. In contrast, our architecture jointly fuses a multi-kernel convolution layer with self-attention within a single sequence block. Multi-size adaptive kernels first learn spatially salient patterns in a dynamic, data-driven manner, which are subsequently globally prioritized through attention. This unified design enables the network to capture both local shape motifs and long-range sequence dependencies simultaneously–a capability not explored in previous work. This property is particularly valuable in the context of GO prediction, where function is often encoded not only in short, localized motifs (e.g., catalytic triads or signal peptides) but also in the coordinated presence of distant sequence elements–such as non-contiguous domain pairs, coevolving residues, or regulatory regions–that interact through long-range dependencies. By jointly modeling both local structural features and global sequence context, the network can effectively associate spatially dispersed yet functionally coupled residues with specific GO terms. In practice, this leads to more accurate predictions for proteins whose functions arise from the interplay of multiple, non-adjacent motifs (e.g., multi-domain enzymes or allosteric receptors), a challenge for traditional motif-based or window-based classifiers.

### Our Contribution

In this work, we present **MkAtt-SDN2GO**, an advanced framework for human protein function prediction that substantially extends the multi-modal data integration principles established by models such as SDN2GO Cai et al. [2020]. Addressing the limitations of previous architectures in capturing the importance of diverse biological features (see sequence module in Figure 1), we introduce a learnable multi-kernel convolutional layer that adaptively extracts local sequence patterns at multiple spatial scales. This is followed by a self-attention mechanism, inspired by Transformer architectures Vaswani et al. [2017], to model long-range dependencies within protein sequence embeddings derived from ProtBERT Elnaggar et al. [2021], enabling precise identification and prioritization of functionally relevant amino acid regions.

- We propose a novel deep learning model that integrates learnable multi-kernel convolution and self-attention mechanisms for accurate protein function prediction.
- We introduce a cross-attention framework to seamlessly integrate protein sequences, domain features, and PPI network embeddings, providing a significant enhancement over the predecessor model.
- Our approach achieves substantial improvements over SDN2GO, including a 14.8% increase in F-max and an 18.8% increase in Recall-max for MF prediction, and a significant F_max_ rise from contemporary comparisons.

**Fig. 1.**
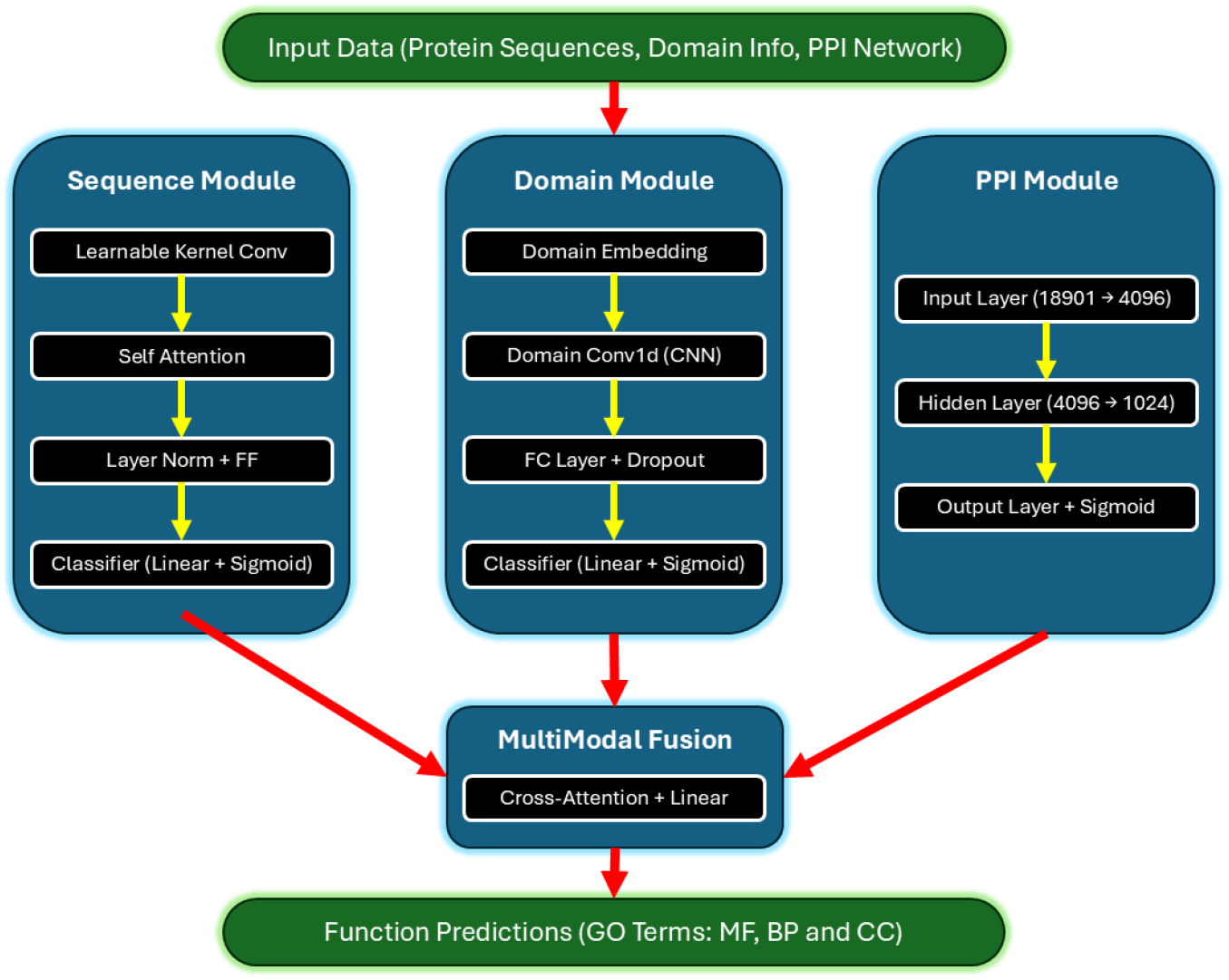
Architecture of the MkAtt-SDN2GO, a GO prediction model. The model comprises three parallel processing streams: (1) Sequence Module with multi-size kernel convolution (referred to as learnable kernel conv) and self-attention for sequence processing, (2) Domain Module with domain embeddings and specialized CNN layers, and (3) PPI Module with dimensionality reduction layers. These streams are integrated through a crossattention based multi-modal fusion mechanism, followed by separate output heads for predicting GO terms across the three ontologies (MF, BP, CC).

## Materials and Methods

### Data Preparation

The construction of the PPI network, multi-scale clustering, and functional annotation via GO enrichment analysis closely follow the methodology established in SDN2GO Cai et al. [2020]. Briefly, high-confidence protein-protein interaction data for uncharacterized human proteins were collected and processed with stringent filtering and topological analysis. Functional modules were identified using MCL and Louvain clustering, followed by GO enrichment analysis (topGO) and semantic summarization (REVIGO); further procedural details are available in the original SDN2GO paper. Some of these details are rewritten here for better readability.

### PPI Network Construction

The construction of the protein-protein interaction network followed a systematic multi-stage process designed to ensure high data quality and relevance. The initial step involved retrieving a list of 8,423 uncharacterized human proteins from the UniProtKB database (release 2023 04). Subsequently, highconfidence interaction data pertaining to these proteins and their interactors were gathered from the STRING database (v11.5) Szklarczyk et al. [2021] and BioGRID (v4.4.229) Oughtred et al. [2021]. The raw network data underwent a preprocessing phase, which included the removal of isolated nodes lacking any interactions and the application of a stringent confidence threshold, retaining only interactions from STRING with a combined score of 400 or greater, and physical interactions from BioGRID. This filtering step resulted in a final network comprising 15,328 protein nodes and 298,765 interaction edges. Finally, a comprehensive topological analysis was conducted using the igraph package (v1.5.1) in R (v4.3.1). This analysis quantified key network properties, including degree distributions, clustering coefficients, and various centrality measures (e.g., degree, betweenness, closeness), to thoroughly characterize the architecture of the resulting PPI network.

### Multi-Scale Clustering

To identify functional modules at different organizational scales within the PPI network, we employed a multi-resolution clustering approach that combined two complementary algorithms. The MCL algorithm Van Dongen [2000], implemented via the ‘MCL’ R package (v1.0.0), configured with an inflation parameter of 1.8, was utilized to reveal fine-grained protein complexes characterized by dense local connections. Concurrently, the Louvain community detection algorithm Blondel et al. [2008], implemented in the igraph package, was applied to identify broader functional modules representing larger biological systems or pathways. The results from both methods were subjected to a comparative analysis for cluster validation, assessing overlap using metrics like the Jaccard index, which revealed a 78% concordance in module assignments between the two techniques. The identified clusters were then filtered to retain only those containing a minimum of five proteins, ensuring biological relevance and facilitating robust downstream enrichment analysis. Following this, a comprehensive characterization of the final clusters was performed, focusing on their size distributions and compositional features (e.g., proportion of characterized vs. uncharacterized proteins).

### Functional Annotation via GO Enrichment Analysis

Functional annotation of the identified protein clusters was achieved through GO enrichment analysis Ashburner et al. [2000], Consortium et al. [2021]. For each cluster identified in the previous step (from both MCL and Louvain), a corresponding gene list (using UniProt IDs) was prepared. Concurrently, comprehensive GO annotations for human proteins (Homo sapiens) were retrieved from the Gene Ontology Consortium. The core of the analysis involved performing GO enrichment using the topGO package (v2.52.0) Alexa et al. [2006] available in R. Fisher’s exact test was applied, incorporating the ‘weight01’ algorithm to account for the GO hierarchy, to statistically assess the overrepresentation of specific GO terms within each cluster compared to the genomic background (all proteins in the constructed network). GO terms were considered significantly enriched if their associated p-value, after applying Benjamini-Hochberg correction for multiple testing, was less than 0.05. The enrichment patterns were analyzed across the three main GO ontologies: MF, BP, and CC. To facilitate interpretation, the enrichment results were visualized using tools like REVIGO Supek et al. [2011] to summarize redundant GO terms based on semantic similarity, highlighting the key functional themes within and across clusters.

### Model Architecture

The model (illustrated in Fig. 1) comprises three parallel processing streams designed to handle sequence, domain, and PPI network data, forming an updated version of SDN2GO with a multi-modal fusion mechanism. Each stream independently encodes its respective input modality prior to integration, and the fused representation supports domain-specific optimization while leveraging cross-domain information.

#### Sequence Module

The protein sequence processing pipeline begins with a learnable multi-kernel convolution block blue(kernel sizes *{*3, 5, 7*}*; shared input from ProtBERT residue embeddings) that adaptively combines outputs from multiple convolution paths. Each path performs a 1D convolution with its respective kernel, and the outputs are combined via a trainable gating vector normalized through a softmax function, enabling the network to assign dynamic importance to different receptive fields. This mechanism is implemented as a ‘LearnableKernelConv’ layer, where the weighted sum of the convolution outputs effectively forms an adaptive convolution that learns which kernel sizes are most relevant for a given sequence. This design enables the model to capture both short- and long-range local motifs without relying on a fixed kernel size.

Following this, a self-attention mechanism (4 heads, dropout 0.1) models long-range dependencies within the sequence, allowing the network to capture contextual relationships between distant residues. The attention sublayer follows a standard Transformer-style configuration with residual connections and layer normalization, where the attention output is passed through a two-layer feed-forward block with ReLU activation and dropout. The module concludes with global average pooling and a linear classifier with sigmoid activation to produce sequence-based function probabilities.

#### Domain Module

Similar to the predecessor model, domain information is first embedded into a continuous vector space. These embeddings are processed through a DomainConv1D layer (a specialized 1D CNN) to capture domain-specific features. The processed features are then passed through a fully connected layer with dropout regularization and sigmoid activation to yield domain-level predictions.

#### PPI Module

Similar to the predecessor model, the PPI network processing stream consists of three main layers: an input layer that reduces the dimensionality from 18,901 to 4,096 features, a hidden layer that further compresses the representation to 1,024 dimensions, and an output layer with sigmoid activation for final networkbased predictions.

#### Multi-Modal Fusion

The outputs from all three modules are integrated through a cross-attention mechanism followed by a linear fusion layer. This enables the model to dynamically weight the importance of each modality based on its relevance for the given functional prediction. In practice, three attention heads (dropout 0.2) are used to compute crossmodal alignment between sequence, domain, and PPI features, enabling the network to capture inter-modal dependencies such as co-evolutionary sequence features correlated with network neighborhood signatures. The fused representation is then forwarded to three ontology-specific output heads corresponding to the MF, BP, and CC tasks.

Overall, the architecture combines adaptive local motif extraction (via learnable multi-kernel convolution) with contextual reasoning (via self- and cross-attention), allowing MkAtt-SDN2GO to jointly exploit sequence composition, domain structure, and network topology within a unified end-to-end trainable framework.

### Evaluation metrics

Following the Critical Assessment of Functional Annotation (CAFA) protocol Zhou et al. [2019], we assessed performance with several complementary measures. Protein-centric precision, recall, and F1-score were computed for each protein after thresholding prediction scores, then averaged across the test set. Term-centric precision, recall, and F1-score were calculated for each GO term and aggregated using both macro-averaging and frequency-weighted micro-averaging. The principal statistic, F-max, is defined as the maximum protein-centric F1 observed when varying the decision threshold from 0 to 1. Threshold-independent assessment was provided by the area under the precision–recall curve (AUPR) and the area under the ROC curve (AUC). Hierarchical precision, recall, and F1-score, which account for parent-child relationships within the GO structure, were also reported to provide a nuanced view of prediction accuracy.

## Results and Discussion

To rigorously evaluate the performance and establish the competitiveness of our integrated functional annotation approach, we conducted extensive benchmarking experiments using 5-fold cross-validation on human protein datasets. The evaluation spanned the three primary GO domains: MF, BP, and CC. The aggregated performance metrics across the folds are summarized in Table 1. Our analysis considered models based on sequence information alone, domain annotations, PPI network structure, and the integrated ‘Weighted’ model, which combines these features. Most of the results of the ‘Domain’ model are higher than the ‘Weighted model’. This behavior was also observed in the original SDN2GO framework. The objective of the weighted (fused) model is not necessarily to outperform every individual modality but to integrate complementary information from the sequence, domain, and PPI features to achieve more balanced and robust predictions across different functional categories. The cross-attention mechanism enables this integration by allowing the model to selectively focus on the most informative aspects of each modality, thereby improving generalization and interpretability.

**Table 1.** Cross-validation test performance of MkAtt–SDN2GO on the Human GO domains. Values are mean *±* standard deviation.

| Model | F-max | AUC | AUPR | Recall-max | Precision-max |
| --- | --- | --- | --- | --- | --- |
| *MF* |  |  |  |  |  |
| Sequence | 0.426 ± 0.005 | 0.922 ± 0.003 | 0.403 ± 0.007 | 0.385 ± 0.016 | 0.476 ± 0.020 |
| Domain | 0.686 ± 0.005 | 0.947 ± 0.005 | 0.693 ± 0.007 | 0.619 ± 0.008 | 0.769 ± 0.006 |
| PPI | 0.356 ± 0.006 | 0.878 ± 0.004 | 0.299 ± 0.005 | 0.312 ± 0.010 | 0.415 ± 0.009 |
| **Weighted** | **0.657 ± 0.003** | **0.918 ± 0.007** | **0.620 ± 0.010** | **0.579 ± 0.012** | **0.761 ± 0.014** |
| *BP* |  |  |  |  |  |
| Sequence | 0.227 ± 0.004 | 0.825 ± 0.004 | 0.175 ± 0.004 | 0.216 ± 0.018 | 0.241 ± 0.011 |
| Domain | 0.438 ± 0.006 | 0.867 ± 0.006 | 0.396 ± 0.006 | 0.349 ± 0.005 | 0.585 ± 0.008 |
| PPI | 0.268 ± 0.005 | 0.816 ± 0.001 | 0.223 ± 0.003 | 0.236 ± 0.008 | 0.309 ± 0.002 |
| **Weighted** | **0.441 ± 0.009** | **0.863 ± 0.004** | **0.384 ± 0.008** | **0.345 ± 0.005** | **0.611 ± 0.020** |
| *CC* |  |  |  |  |  |
| Sequence | 0.473 ± 0.006 | 0.923 ± 0.003 | 0.467 ± 0.007 | 0.428 ± 0.015 | 0.528 ± 0.016 |
| Domain | 0.519 ± 0.011 | 0.920 ± 0.012 | 0.516 ± 0.020 | 0.479 ± 0.024 | 0.569 ± 0.018 |
| PPI | 0.441 ± 0.004 | 0.895 ± 0.003 | 0.395 ± 0.010 | 0.469 ± 0.009 | 0.416 ± 0.005 |
| **Weighted** | **0.522 ± 0.013** | **0.906 ± 0.011** | **0.492 ± 0.023** | **0.452 ± 0.012** | **0.617 ± 0.018** |

### Comparative Analysis

To accurately position our findings within the current landscape of protein function prediction, we performed a critical comparison of our model against some of the contemporary methods. Figure 2 contrasts our MkAtt-SDN2GO results with representative baselines reported for the *Homo sapiens* species in previous studies, including GOHPro and several other established methods Hu and Zhao [2025]. Although the underlying ontology categories (BP, MF, and CC) and evaluation metrics remain consistent, the dataset usage and experimental settings vary slightly across studies. Hence, the presented plots are intended to provide an indicative comparison of overall performance trends on the same species, rather than a direct one-to-one benchmarking under identical conditions.

**Fig. 2.**
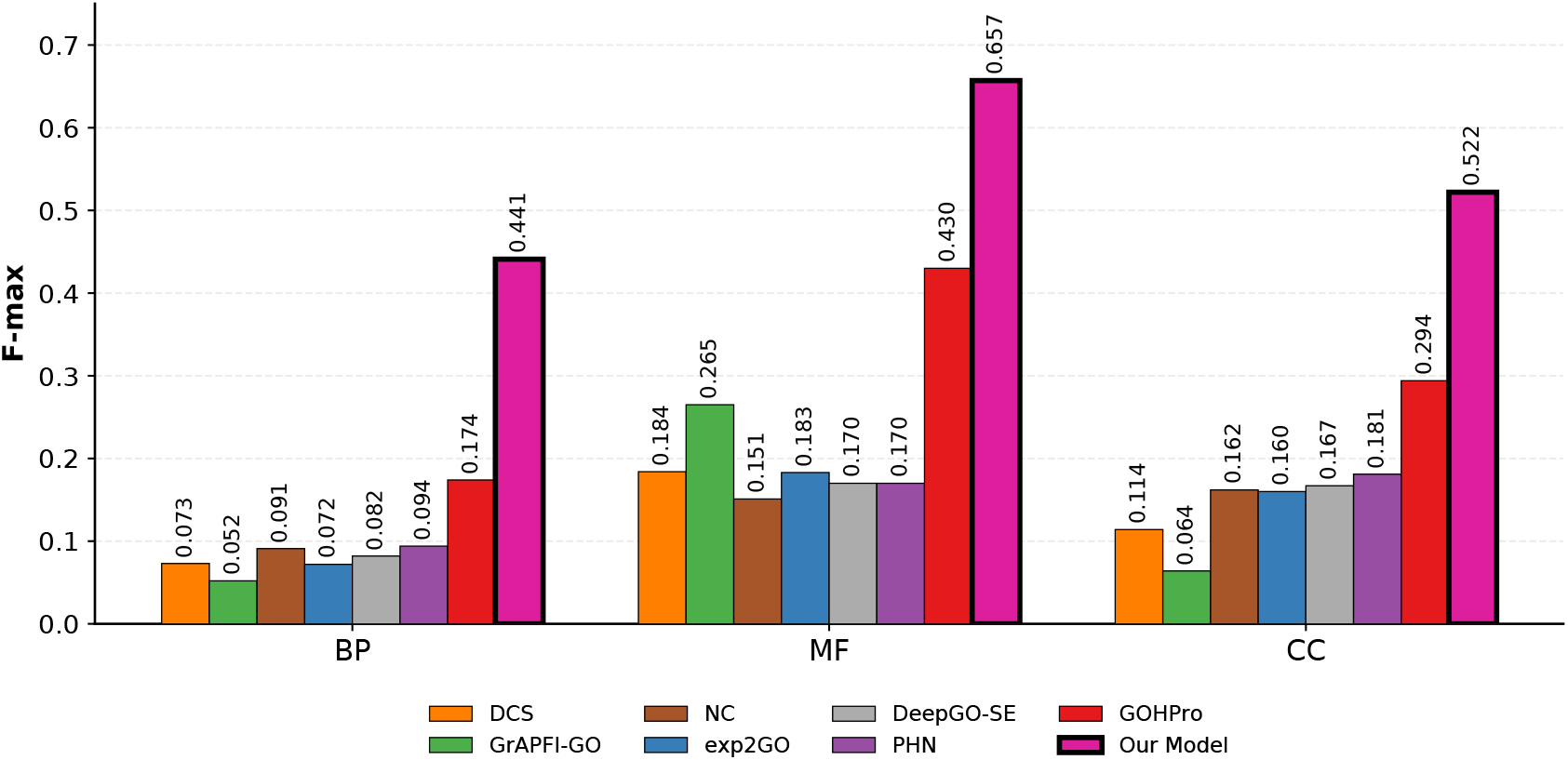
F-max comparison across GO domains (*Homo sapiens*). The figure compares contemporary methods such as GOHPro, exp2GO, GrAPFI-GO, PHN, DCS, NC, and DeepGO-SE. The results correspond to the BP, MF, and CC domains for the human species. The comparison follows the standard CAFA evaluation framework based on the F-max metric.

The comparative advantage is particularly evident when examining the Fmax metrics, where our approach shows consistent improvements over these comparison methods. This performance advantage can be attributed to our model’s ability to effectively integrate multiple data modalities and leverage the attention mechanism to focus on the most informative features for each prediction task.

Within this context, our model achieves the top *F* -max in each ontology: 0.441 (BP), 0.657 (MF), and 0.522 (CC). Relative to the strongest baseline reported in Hu and Zhao [2025], these correspond to absolute gains of +0.267 in BP (vs. GOHPro, 0.174), +0.227 in MF (vs. GOHPro, 0.430), and +0.228 in CC (vs. GOHPro, 0.294).

Compared to (i) network-only or network-dominant methods (PHN, DCS, NC), which can underperform when PPI coverage is sparse or noisy; (ii) expression-driven exp2GO, which depends on compendia that may lack breadth for many proteins or tissues; (iii) semantic/graph methods (GrAPFI-GO, DeepGO-SE) that chiefly exploit ontology structure and textual/semantic relations; and (iv) propagation in GOHPro that relies on well-annotated neighborhoods, our model explicitly fuses three complementary evidence streams (sequence, domain, PPI) with a learnable multi-kernel sequence encoder and cross-attention for modality weighting. The multi-kernel block discovers motif-scale patterns without fixing a receptive field, while cross-attention prioritizes the most informative modality per term, mitigating failure modes where any single source is weak. This design yields robust protein-centric discrimination under CAFA scoring, particularly on MF and CC where sequence motifs and local cellular context are jointly informative.

Compared to traditional methods relying heavily on sequence similarity (e.g., BLAST) or homology transfer, our approach can annotate proteins lacking close homologs by leveraging network context and learned sequence patterns. While structural prediction methods provide valuable functional clues, they often require significant computational resources and may not be feasible for all proteins; our method integrates available sequence and network data efficiently. Compared to methods based solely on gene expression correlation, which infer functional links from co-expression patterns, our PPI-based approach captures direct physical interactions, providing a different, often complementary, layer of functional evidence. Within the realm of machine learning, our attention-based model surpasses many traditional algorithms and simpler deep learning architectures (e.g., sequence-only CNNs or RNNs) by explicitly modeling both sequence features and network topology through attention mechanisms. A key differentiator is the interpretability offered by attention weights, which provides insights often lacking in ‘black-box’ prediction models.

### Comparison with SDN2GO Baseline

Given that our architecture builds upon and enhances the SDN2GO framework, a direct comparison with SDN2GO’s performance is particularly relevant. We followed the 5 fold cross validation protocol consistent with SDN2GO Table 1 and observed substantial improvements across all key metrics in the MF domain, as shown in Figure 3. We reimplemented SDN2GO under our evaluation setup and observed a discrepancy in the reproduced scores compared to those reported in its Table 1, which may be due to differences in random initialization and the fact that their cross-validation was performed on an 80% training split, with a separate 20% held out for final testing. Our experiments did not include such an independent test set.

**Fig. 3.**
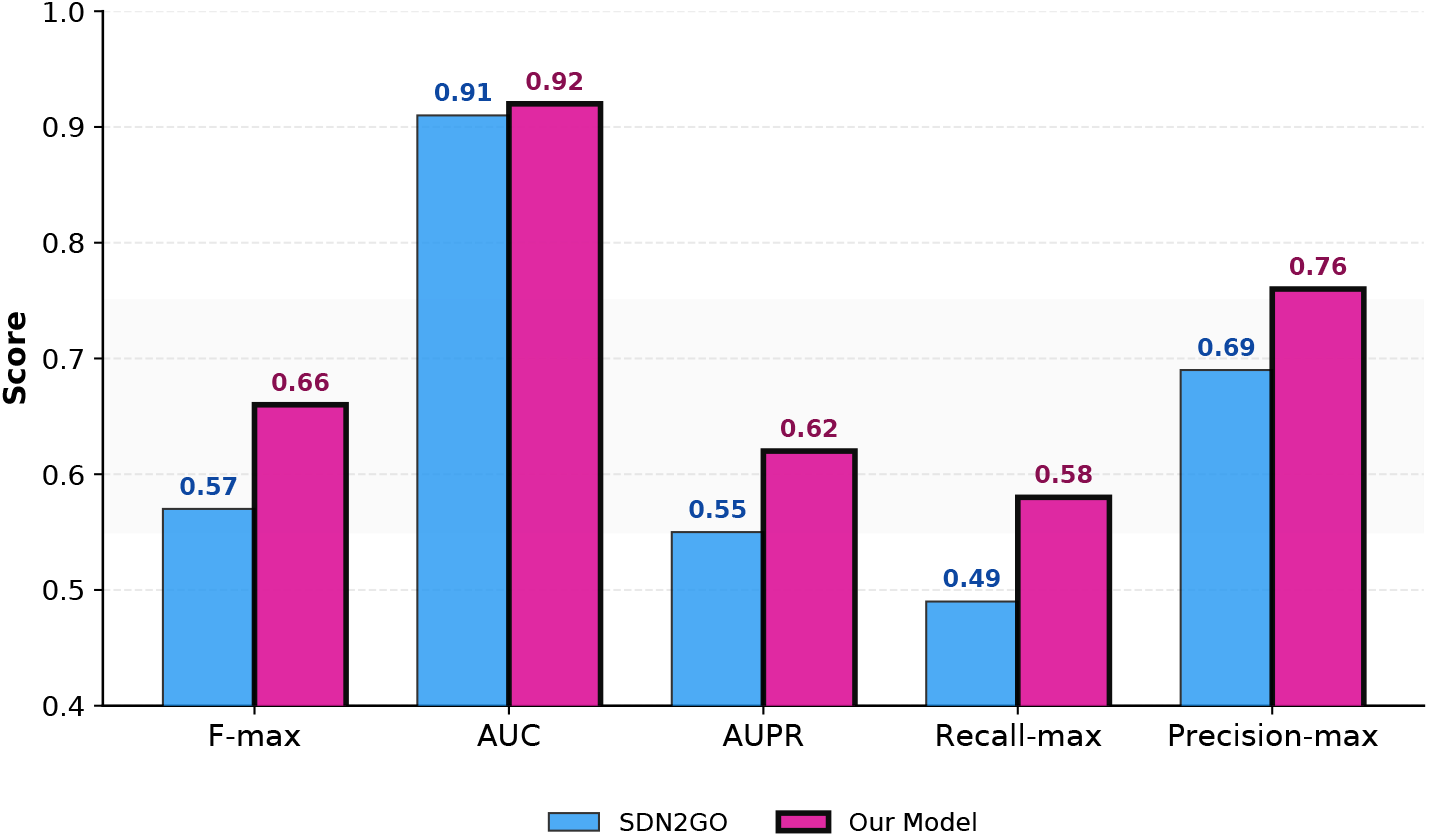
Detailed comparison with SDN2GO baseline for MF prediction. Our Model shows consistent improvements across all metrics, with particularly notable gains in F-max (14.8% improvement) and AUPR (14.1% improvement).

Most notably, our weighted model achieves an F-max score of 0.657 compared to SDN2GO’s 0.572, representing a 14.8% improvement. Similar enhancements are observed across other metrics, with our model showing a 14.1% increase in AUPR (0.620 vs 0.543), an 18.8% improvement in Recall-max (0.579 vs 0.487), and a 9.8% gain in Precision-max (0.761 vs 0.693). The AUC score also shows a modest improvement of 1.2% (0.918 vs 0.907), indicating better overall discriminative capability.

These improvements can be attributed to several key architectural enhancements in our model, particularly for the sequence module. The introduction of attention mechanisms, particularly in processing sequence and network features, allows for more nuanced feature extraction compared to SDN2GO’s CNN-based approach. Our model’s ability to dynamically weight different aspects of the input data using the multi-size learnable kernels, rather than relying on static feature extraction, appears to be especially beneficial for capturing the complex patterns necessary for accurate MF prediction. Furthermore, the improved performance metrics suggest that cross-attention-based dynamic weighing better handles the inherent complexity of protein function prediction, particularly in identifying specific molecular activities and binding patterns characteristic of the MF domain.

### Effectiveness of the Integrated Approach

The results presented herein strongly support the effectiveness of integrating network-based clustering with attention-based deep learning for protein function annotation. This synergy leverages the complementary strengths of both methodologies. The network-based clustering component excels at identifying functional modules and placing uncharacterized proteins within a broader biological context based on interaction patterns (‘guilt by association’). This approach effectively groups proteins that cooperate in specific cellular processes. Concurrently, the attention-based deep learning model, particularly the self-attention, captures intricate sequence-specific features and complex, weighted relationships within the interaction network that contribute directly to a protein’s function. By combining these perspectives-the community context from clustering and the feature-specific insights from the deep learning model, we achieve functional annotations that are comprehensive, robust, and confident. The interpretability afforded by the attention mechanism further enhances the approach, providing plausible biological rationales for the functional predictions by highlighting salient sequence motifs and influential interaction partners. The observation that consensus annotations derived from both methods exhibit superior precision and recall underscores the value of this integrative strategy in reducing prediction errors and increasing reliability.

### Limitations

While the integrated approach demonstrates considerable promise, it is essential to acknowledge certain limitations inherent in the methodologies and data sources employed. The accuracy and completeness of the underlying PPI data are critical; both false positive interactions (spurious links) and false negative interactions (missing true links) within databases like StringDB can impact network structure and subsequent analyses, potentially leading to incorrect clustering or biased network embeddings. Consequently, our analysis is inherently limited to proteins for which interaction data is available, potentially excluding proteins that function in isolation or whose interactions are poorly characterized. It is crucial to reiterate that the functional annotations generated are computational predictions and require experimental validation to be definitively confirmed. The current models primarily capture static interaction networks and may not fully represent the dynamic nature of protein interactions, which can vary significantly depending on cellular conditions, developmental stage, or environmental stimuli. The performance of the attention-based model, while strong overall, exhibited variability across different GO categories, with predictions for very specific or sparsely annotated GO terms remaining challenging due to limited training examples. Addressing these limitations represents important directions for future refinement.

Further development could focus on designing more sophisticated attention mechanisms, perhaps hierarchical attention models that explicitly mirror the structure of protein domains or graph transformer architectures capable of capturing longer-range dependencies within the PPI network. Applying and rigorously benchmarking the integrated pipeline on proteomes from diverse organisms, beyond yeast and human, would assess its generalizability and potentially reveal conserved functional modules. A crucial next step involves the experimental validation of high-confidence predictions made for selected uncharacterized proteins; collaboration with experimental biologists would be invaluable for designing and executing these validation studies. Methodologically, extending the model to predict more specific, lower-level GO terms and to improve performance on rare functional categories remains an important challenge, potentially requiring techniques for handling extreme class imbalance or leveraging the GO hierarchy more explicitly during training or inference. Finally, developing a user-friendly web-based tool or software package implementing this functional annotation pipeline would greatly benefit the broader research community, enabling wider adoption and application of these powerful integrative methods.

## Conclusion

We introduced **MkAtt–SDN2GO**, a human protein function prediction model that couples a *learnable multi-kernel* sequence encoder with *self-attention* and a *cross-attention* fusion of sequence, domain and PPI signals. On CAFA-style, time-delayed evaluation, our Weighted (fused) model achieves comparable protein-centric performance for *Homo sapiens*, with top F_max_ across all ontologies (BP: 0.441, MF: 0.657, CC: 0.522) and consistent gains over SDN2GO on MF (e.g., +14.8% in F_max_). A comparison with representative human-species baselines compiled from the literature further indicates clear advantages for our approach under the same metric.

The *multi-kernel* block adapts its receptive field to the local motif scale, allowing the sequence stream to discover short signatures (e.g., catalytic/ligand motifs) and longer patterns without fixing a priori window sizes, but making it learnable based on multi-sized kernels usage. *Self-attention* then aggregates long-range residue dependencies that often underlie function (e.g., non-contiguous domain cooperativity). Finally, *cross-attention* acts as a learned gate over modalities, up-weighting the most informative source term-by-term. This threefold design-adaptive motif capture, global sequence reasoning, and modality re-weighting, motivates the name *Multi-kernel Attentive*.

In several tasks, our *Domain* stream alone is very strong and can occasionally edge the *Weighted* fusion in cross-validation means. This behavior mirrors SDN2GO and reflects the fact that InterPro-like domain signals are highly discriminative for many MF/CC terms. Nevertheless, the cross-attention fusion is important: (i) it improves robustness when domain evidence is weak or missing, (ii) it yields a better precision–recall trade-off on average, (iii) it increases coverage by leveraging sequence and network cues for proteins with sparse domain annotations, and (iv) it adds interpretability—attention weights expose which modality, region or neighbor drove a prediction.

Relative to baselines collated from Hu and Zhao (GOHPro) and related methods for human proteins, our model attains a comparable protein-centric F_max_ in BP, MF and CC (Fig. 2). Methodologically, MkAtt–SDN2GO differs by unifying (i) adaptive motif discovery (multi-kernel CNN), global sequence reasoning (self-attention), and (iii) data-driven modality selection (cross-attention). This combination mitigates failure modes of single-evidence systems: propagation methods degrade with sparse neighborhoods; expression-driven models depend on compendia breadth; network-only models suffer under PPI incompleteness; and semantic/graph approaches alone underutilize raw sequence signal.

We follow the CAFA protocol (protein-centric F_max_, time-delayed assessment) because it standardizes evaluation across ontologies and enables comparison to widely reported human-species numbers. While CAFA5 has recently emerged, comparable public, *human-species* breakdowns for all reference methods used in Fig. 2 are not uniformly available. We therefore prioritize a like-for-like species/matrix comparison in the CAFA2/3 style to avoid confounding changes in ontology versions, evidence codes, or leaderboard composition. The proposed architecture is orthogonal to the challenge edition and can be evaluated on CAFA5 as those per-species, per-ontology statistics become accessible. This can be addressed in the future.

Performance still varies across sparsely annotated terms; PPI incompleteness and static networks remain bottlenecks. Future extensions include: integrating structure and expression as additional modalities within the same cross-attention fusion; hierarchy-aware training to better exploit GO ancestry; calibrated uncertainty for experimental prioritization; and broader organismal evaluation, including prospective CAFA5-formatted assessments as standardized human-species baselines solidify.

MkAtt–SDN2GO demonstrates that *adaptive, attentive sequence modeling* combined with *learned, interpretable multi-modal fusion* yields reliable human protein function predictions under CAFA-style scoring. Even where a domain stream is dominant, cross-attention improves robustness, coverage, and insight—key properties for practical largescale annotation. We release code and artifacts to facilitate reproduction and downstream use.

## List of Abbreviations

GO: Gene Ontology
MF: Molecular Function
BP: Biological Process
CC: Cellular Component
PPI: Protein-Protein Interaction
CNN: Convolutional Neural Network
GCN: Graph Convolutional Network
SDN2GO: Shape-aware Deep Network for Gene Ontology prediction (baseline model)
F-max: Maximum F1 Score
Recall-max: Maximum Recall

## Declarations

### Availability of data and materials

The datasets used in this study were derived from publicly available biological databases, including StringDB, BioGRID, and UniProtKB. These databases are accessible at:

- https://string-db.org/
- https://thebiogrid.org/
- https://www.uniprot.org/

Additional data and intermediate results produced using opensource R packages (e.g., `igraph, MCL, topGO, GOSemSim`) and Python libraries (e.g., `PyTorch, PecanPy, ProtBERT`) are available from the corresponding author upon reasonable request.

### Authors’ contributions

K.J. and R.S. conceived the main idea. T.B. performed the literature survey. K.J. performed model development, experiments, analysis, and interpretation of the results. K.J., T.B., and W.L. contributed to manuscript writing and revision.

## Acknowledgements

This research is supported by the Ministry of Education, Singapore, under its Research Centre of Excellence award to the Institute for Digital Molecular Analytics & Science, NTU (IDMxS, grant: EDUNC-33-18-279-V12).

## Code availability

All source code supporting this study can be made available upon reasonable request to the corresponding author.

